# A Core Pattern of Cerebellar and Brainstem Degeneration and Reduced Cerebrocerebellar Structural Covariance in Spinocerebellar Ataxia Type 3 (SCA3): MRI Volumetrics from ENIGMA-Ataxia

**DOI:** 10.1101/2025.08.08.669370

**Authors:** Jason W. Robertson, Isaac Adanyeguh, David J. Arpin, Tetsuo Ashizawa, Benjamin Bender, Fernando Cendes, Xi Chen, Giulia Coarelli, Léo Coutinho, Andreas Deistung, Imis Dogan, Alexandra Durr, Jennifer Faber, Juan Fernandez-Ruiz, Mónica Ferreira, Marcondes C. França, Sophia L. Göricke, Shuo Han, Thomas Klockgether, Chen Liu, Jun Luo, Alberto R. M. Martinez, Sergio E. Ono, Chiadi U. Onyike, Gülin Öz, Henry Paulson, Jerry L. Prince, Kathrin Reetz, Thiago J. R. Rezende, Matthis Synofzik, Hélio A. Ghizoni Teive, Sophia I. Thomopoulos, Paul M. Thompson, Dagmar Timmann, David Vaillancourt, Bart van de Warrenburg, Judith van Gaalen, Xingang Wang, Philipp Wegner, Sarah H. Ying, ESMI MR Study Group, EUROSCA MR Study Group, Ian H. Harding, Carlos R. Hernandez-Castillo

## Abstract

**Objective:** Spinocerebellar ataxia type 3 (SCA3) is a rare, inherited neurodegenerative disease. Here, we profile the spatial spread of atrophy across the whole brain, determine whether brain degeneration preferentially maps onto specific functional networks, and investigate the relationship between cerebellar and cerebral anatomical changes.

**Methods:** Whole-brain grey and white matter (GM and WM) voxel-based morphometry was performed on 408 individuals with SCA3 (82 pre-ataxic) and 293 controls. The SCA3 cohort was stratified by ataxia severity to investigate disease progression, with cerebellar GM atrophy mapped onto a task-based functional atlas. Volume was correlated with disease duration and intensity. Cerebrocerebellar volumetric covariance was assessed to determine whether atrophy was coupled between infra- and supratentorial regions.

**Results:** The pattern of atrophy is spatially consistent but progressive in magnitude across the disease course. The greatest atrophy was found in the pons, cerebellar WM, and cerebellar peduncles; correlations with disease severity and duration were also strongest in these regions. Cerebellar GM atrophy was greatest in functional regions associated with motor execution and planning, attention, and emotional processing. Sparse cerebral cortical atrophy appears only in the most severe disease subgroup, while striatal atrophy begins in the earliest stages. Reduced cerebrocerebellar structural covariance is observed in SCA3 participants versus controls.

**Interpretation:** While cerebellar and brainstem atrophy become more severe, the pattern of atrophy remains largely consistent as SCA3 progresses. Cerebellar GM degeneration occurs in regions associated with motor, cognitive, and affective control, in line with clinical presentation. Cerebellar atrophy is not directly mirrored by cerebral changes.

## Introduction

Spinocerebellar ataxia type 3 (SCA3), also known as Machado-Joseph Disease, is a neurodegenerative disease caused by a CAG triplet repeat expansion in the ATXN3 gene on the long arm of Chromosome 14^1^, resulting in misfolding and aggregation of the ataxin-3 protein.^2^ This in turn results in neuronal cell death and/or demyelination in the spinal cord, cerebellum, brainstem, striatum, and thalamus, as well as in the tracts that link these structures.^1–3^ SCA3 is defined by progressive cerebellar ataxia, and variable expression of neurological symptoms including peripheral neuropathy, bradykinesia, tremor, Parkinsonism, oculomotor impairment, and dysphagia.^1,4,5^

Non-invasive imaging techniques such as magnetic resonance imaging (MRI) have been crucial to describing the neuroanatomical changes underlying SCA3 and their relationship with clinical disease severity. A growing body of MRI research has examined the progression of SCA3 both longitudinally^6–11^ and cross-sectionally.^9,12–15^ This research indicates that regional atrophy is progressive in both spatial extent and severity, including an evolving pattern of infratentorial-to-cerebral atrophy^13^ and distinct time courses of volume loss in cerebellar, brainstem, and striatal structures.^12^ The extent of atrophy in key brain regions is correlated with motor impairment severity^11,12,14,16^, as well as cognitive impairment consistent with cerebellar cognitive affective syndrome (CCAS).^14,17–19^

Understanding the development of SCA3 in both time and space is critical for predicting clinical variability and progression, selecting imaging biomarkers for clinical translation, and targeting treatments. While many studies have undertaken these tasks by examining large anatomical regions of interest (ROIs)^11,12,15,20,21^, voxel-based approaches can provide greater specificity and more spatially nuanced inferences from atrophy patterns.

Here we seek to expand on the existing knowledge base using MRI data from a large, retrospective, multi-site cohort of individuals with SCA3 spanning a broad spectrum of disease severity, including pre-ataxic individuals. Using voxel-based morphometry (VBM), we first seek to define the profile of whole-brain atrophy in participants with SCA3 at different stages of ataxia severity. We then examine how the SCA3 atrophy pattern maps onto a novel functional parcellation of the cerebellum^22^, allowing insights to be derived by mapping atrophy to regions responsible for motor and non-motor functions within the cerebellum. Finally, we evaluate the relationship between cerebellar and cerebral atrophy using corticocerebellar covariance analysis to determine whether there is direct correspondence between volume loss in the cerebrum and cerebellum.

## Methods

### Participants and Data

This study used a retrospective, cross-sectional design with data aggregated through the ENIGMA-Ataxia working group. All procedures were approved by both the Monash University Human Research and Ethics Committee and the Dalhousie University Research Ethics Board. Structural MRI, demographics, and clinical data were collected from participants with SCA3 (*n* = 408) and healthy controls (CONT; *n* = 293) from eleven sites (Aachen, Germany; Baltimore, USA; Campinas, Brazil; Chongqing, China; Curitiba, Brazil; Essen/Halle, Germany; Florida, USA; Mexico City, Mexico; Nijmegen, Netherlands; Paris, France; and Tübingen, Germany) and three collaborating consortia (ESMI, EUROSCA, and READISCA) in accordance with their respective ethics and governance bodies. All data were anonymized and given new subject identifiers prior to aggregation.

Participants with SCA3 were included based on: (1) at least 52 CAG repeats in *ATXN3*^1^, including individuals with and without manifest ataxia, and/or (2) genetically confirmed family history of SCA3 with clinical manifestation of disease-consistent ataxia. Disease duration was recorded as the time since the patient first recognised their ataxia symptoms. Ataxia severity at the time of the scan was quantified using the Scale for Assessment and Rating of Ataxia (SARA)^23^. Subjects were initially excluded on the basis of (1) missing age or inconsistent clinical date, or (2) image quality or other technical issues; after some initial analyses (see Supplemental Results), subsequent exclusions were performed on specific sites due to a lack of control data, or due to small samples within site (Figure 1).

**Figure 1.**
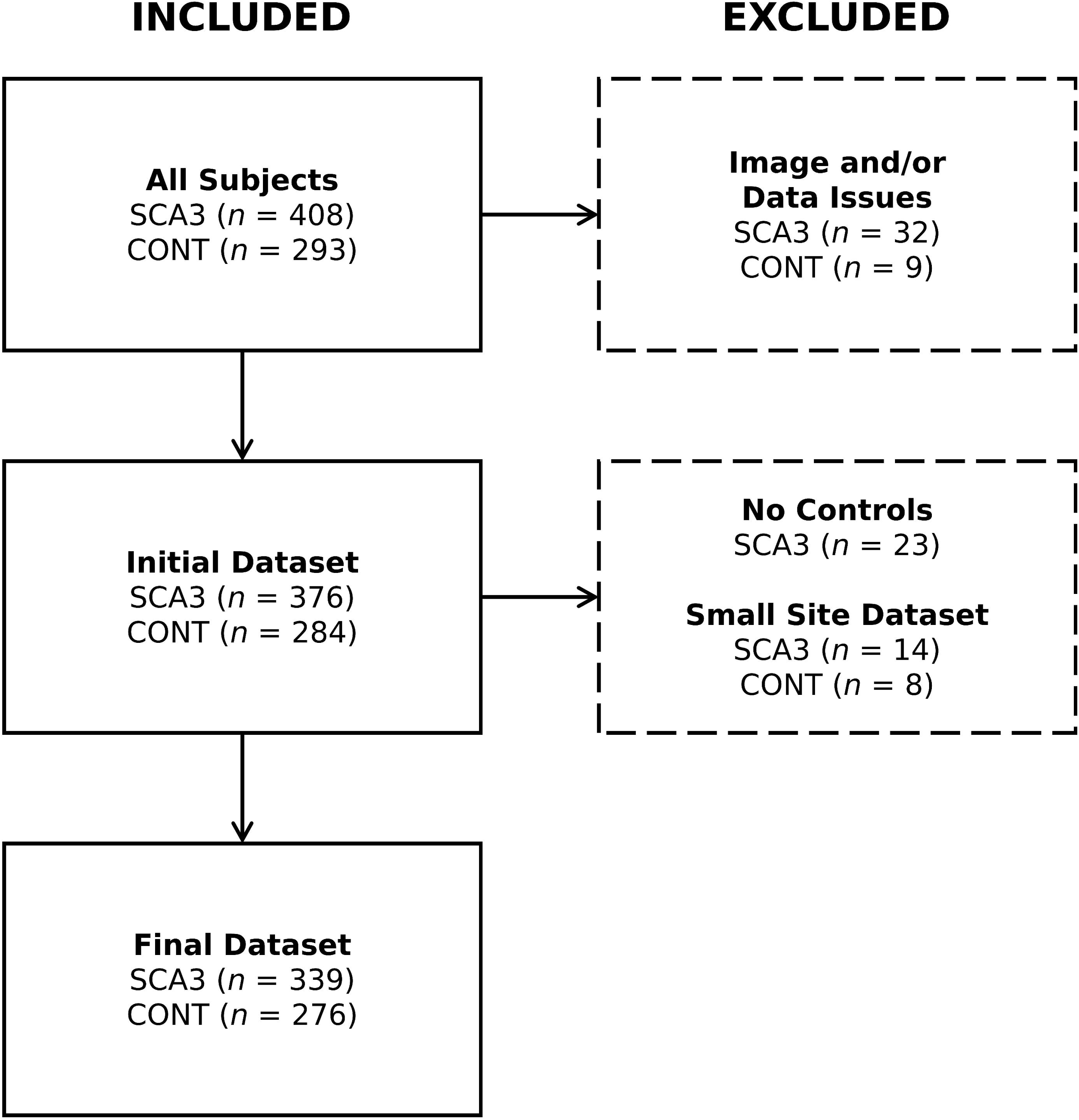
Flowchart illustrating the subject exclusion process.

A whole-brain, high-resolution T_1_ -weighted structural MRI image (voxel size 1 mm^3^ or less) was collected for each subject. Imaging protocols varied by site, but were consistent between SCA3 and CONT cohorts at each site (Supplementary Table S1).

### Image Processing

All T_1_ -weighted images were preprocessed using a previously-described algorithm^24^, which combines specialized software for analyzing the cerebellum and cerebrum.^25^ First, a cerebellar mask was derived for each image using the ACAPULCO^26^ algorithm (v0.2.1). The automated masks were visually inspected, with minor segmentation errors corrected manually and major errors resulting in exclusion. Next, cerebellar grey matter (GM) was extracted using the Spatially Unbiased Infratentorial Toolbox (SUIT)^27^ v3.2 for SPM12^28^ v7771, as implemented for MATLAB (Natick, USA). Using the previously generated cerebellar mask, the images were segmented into GM and white matter (WM) partial volumes, which were DARTEL normalized and resliced into SUIT space; Jacobian modulation was used to ensure that the value assigned to each voxel remained proportional to its original volume. The resulting cerebellar GM images were inspected for normalization errors, and the spatial covariance of these images was calculated and compared across the entire cohort: outliers were visually inspected to determine if rejection was necessary due to processing errors. Finally, the resulting images were spatially smoothed using a Gaussian kernel at a full width at half-maximum (FWHM) of 3 mm.

Because SUIT is optimized for cerebral GM, whole-brain WM and cerebral GM were calculated using the Computational Anatomy Toolbox (CAT12, v12.5)^29^ for SPM12. The raw images were bias-field corrected, skull-stripped then segmented into GM, WM, and cerebrospinal fluid (CSF) partial volumes, from which intracranial volume (ICV) could be calculated. These images were also DARTEL registered into Montreal Neurological Institute (MNI) space with Jacobian modulation to maintain volume encoding. The GM image was masked to remove the cerebellum, to prevent spatial smoothing from causing undue influence on the cerebral results. Finally, both the GM and WM images were smoothed with a 5 mm FWHM Gaussian kernel.

### Statistical Analysis

Demographic and clinical analyses were undertaken using R v4.3.3. The distribution of ages in each group was first checked for normality using the Shapiro-Wilk test; if at least one group was not normally distributed, the Wilcoxon ranked-sum test was used to determine if groupwise differences were significant. Sex was compared between groups using Fisher’s exact test.

All image-based statistical analyses were performed using SPM12 in MATLAB. In all cases, heteroscedasticity was assumed and corrected for, with final voxel-level inferences estimated with family-wise error (FWE) corrections at the *α* = 0.05 level, using random field theory to account for the large number of multiple comparisons endemic to imaging analysis.^30^ Only significant clusters with *k* ≥ 100 voxels were reported, ensuring robust but conservative inference. Statistical maps showing significant results were then converted to Cohen’s *d* (groupwise analyses) or Pearson’s *r* (correlation analyses) to describe effect sizes.^31,32^

#### Between-Group Analysis for Disease Effects

Groupwise comparisons between SCA3 and CONT participants were performed by creating general linear models (GLMs) in SPM12 with Group (2 levels) and Site as factors, and ICV and Age as covariates. For the initial dataset, Site had 14 levels, while for the final dataset (Figure 1), Site had 11 levels. Of these, only Group is a comparison of interest; the remainder are included as nuisance variables.

For interpretative purposes, the voxel-based results were mapped to one of several anatomical atlases: the Harvard-Oxford cortical and subcortical GM atlases and the Johns Hopkins University cortical WM atlas as included with FSL^33^; the SUIT cerebellar grey matter and dentate atlases^34,35^; the van Baarsen cerebellar WM atlas^36^; and the FreeSurfer brainstem atlas.^37^ Significant results in the cerebellar GM were additionally mapped onto a multi-domain task-based (MDTB) parcellation developed by King *et al.*^22^ to determine correspondence between atrophy and functional regions of the cerebellum.

#### Clinical Correlations for SCA3 Participants

In the initial dataset, the spatially smoothed SCA3 images were assessed directly for linear relationships between volume and either disease duration or SARA score; subject Age and ICV were incorporated as covariates of no interest. Subsequently, images for which the effects of healthy aging^38^ and site-specific confounds^39^ could not be estimated and adjusted for^40^ were excluded (Figure 1). The age- and site-adjusted images were then re-analyzed, removing Age but retaining ICV as a covariate of no interest. For both comparisons, estimated time to disease onset was calculated for the pre-symptomatic cohort based on the methodology of Tezenas du Montcel *et al.*^41^ when possible and included as a negative disease duration value.

#### Disease Staging

Cross-sectional analyses were performed on subsets of the SCA3 participant data to track disease progression. The data were subdivided into five groups – pre-symptomatic (SARA < 3)^42^ and four quartiles of symptomatic individuals (quartile separation values: SARA = 8, 11, and 16) – which were then compared to the full CONT cohort using a GLM with ICV as a nuisance covariate. To minimize the possibility of site effects in the data, these analyses were undertaken with the age- and site-corrected images generated for the correlations.

#### Cerebrocerebellar Covariance

To better understand the relationship between cerebellar and cerebral disease effects, we undertook a cerebrocerebellar covariance analysis, using the following linear model:

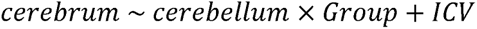

Where *cerebrum* and *cerebellum* represent the respective GM, WM, or total volumes in those regions of the brain, *Group* is a binary variable representing CONT (0) or SCA3 (1), and *ICV* is intracranial volume as a covariate. This model was run for GM only, WM only, and the two summed together. For this analysis only, the SUIT-derived cerebellar WM was processed using the same procedure as the cerebellar GM and used in lieu of the CAT12-derived whole-brain WM, while the CAT12-derived WM was cropped to the cerebrum.

As this analysis was intended to examine the extent to which cerebellar and cerebral atrophy are linked, the Group interaction was the comparison of interest: stronger correlations in the SCA3 cohort versus CONT would imply that atrophy in the cerebellum and cerebrum are linked, while a weaker correlation would imply that the two are relatively independent.

## Results

Demographic and clinical data from the final dataset are summarized in Table 1. The SCA3 and CONT cohorts were age- and sex-matched within each site, and in aggregate showed no significant differences in either sex (*χ*^2^ = 0.921; *p* = 0.383) or age (Wilcoxon *W* = 47360.5, *p* = 0.792).

**Table 1.** Summary of subject demographic and clinical characteristics.

|  | SCA3 |  |  |  |  |  |  | CONT |  |  |
| --- | --- | --- | --- | --- | --- | --- | --- | --- | --- | --- |
| Site | N | F (%) | Age | Disease Duration | CAG Repeats | SARA | Pre-Ataxic (%) | N | F (%) | Age |
| Baltimore | 11 | 8 (72.7) | 51.6 (10.1) | 8.3 (4.5) | 70.3 (3.0) | 10.8 (5.4) | 1 (9.1) | 12 | 8 (66.7) | 48.6 (11.8) |
| Campinas | 79 | 43 (54.4) | 48.1 (12.6) | 10.4 (7.0) | 72.0 (3.7) | 13.5 (8.6) | 5 (6.3) | 82 | 43 (52.4) | 47.5 (11.7) |
| Chongqing | 42 | 18 (42.9) | 41.2 (12.4) | 8.4 (4.1) | 66.2 (4.3) | 10.5 (8.3) | 8 (19.0) | 55 | 27 (49.1) | 41.9 (13.0) |
| Curitiba | 16 | 7 (43.8) | 44.6 (13.2) | 10.8 (7.0) | --<sup>a</sup> | 13.4 (5.5) | 0 (0.0) | 17 | 8 (47.1) | 43.9 (11.9) |
| ESMI | 59 | 30 (50.8) | 48.1 (11.5) | 9.5 (5.7) | 69.2 (4.1) | 9.4 (7.0) | 14 (23.7) | 21 | 10 (47.6) | 50.0 (13.9) |
| Essen/Halle | 22 | 11 (50.0) | 52.0 (13.9) | 13.5 (7.0) | 68.2 (4.1) | 10.1 (5.3) | 3 (13.6) | 21 | 8 (38.1) | 51.8 (14.3) |
| Mexico | 16 | 9 (56.3) | 40.3 (12.3) | 6.9 (4.7) | 72.6 (3.9) | 13.3 (7.7) | 2 (12.5) | 15 | 7 (46.7) | 41.7 (13.3) |
| Nijmegen | 17 | 8 (47.1) | 37.4 (9.6) | --<sup>b</sup> | 66.8 (2.9) | 1.5 (1.0) | 17 (100.0) | 8 | 2 (25.0) | 31.3 (8.2) |
| Paris | 20 | 12 (60.0) | 50.9 (11.8) | 8.7 (4.5) | 68.9 (5.6) | 13.0 (7.2) | 3 (15.0) | 24 | 13 (54.2) | 49.8 (12.8) |
| READISCA | 48 | 30 (62.5) | 43.8 (10.1) | 4.8 (4.4) | 70.4 (3.6) | 4.2 (3.1) | 19 (39.6) | 12 | 6 (50.0) | 41.3 (7.5) |
| Tübingen | 9 | 3 (33.3) | 40.4 (8.3) | 9.0 (5.7) | 67.6 (2.4) | 3.3 (4.0) | 6 (66.7) | 9 | 3 (33.3) | 41.0 (9.0) |
| **Total** | **339** | **179 (52.8)** | **45.9 (12.3)** | **9.2 (6.1)** | **69.6 (4.4)** | **9.9 (7.7)** | **78 (23.0)** | **276** | **135 (48.9)** | **45.6 (12.8)** |

Parentheses indicate standard deviation except where indicated. Age and disease duration are measured in years. N: number of subjects; F: number of female subjects; SARA: Scale for the Rating and Assessment of Ataxia score.
<sup>a</sup> Data not available.
<sup>b</sup> No manifest ataxia subjects.

### Volumetric Differences Between SCA3 Participants and Healthy Controls

The GLM comparing SCA3 and CONT participants found numerous areas of significant atrophy. Those with the greatest effect sizes were observed in the WM of the brainstem and cerebellum (*d* > 1.5; Figure 2A), including the medial lemniscus, dentate region, pons, pontine crossing tract, corticospinal tract, and the cerebellar peduncles. Significant but less robust atrophy (0.4 < *d* < 1.5) was observed in the midbrain, medulla, and internal capsule. In the cerebellar GM, atrophy at a medium to large effect size was evident in all regions (*d* < 1.0; Figure 2B), most notably in bilateral Lobule VI, Vermis VIIb and IX, and right Lobule IX. In the cerebral GM (Figure 2C), only the bilateral caudate nuclei and dorsal-anterior putamen were significantly atrophied in SCA3 participants compared to controls. A small area of the bilateral thalamus was found to have greater volume in SCA3 participants (Supplementary Figure S1).

**Figure 2.**
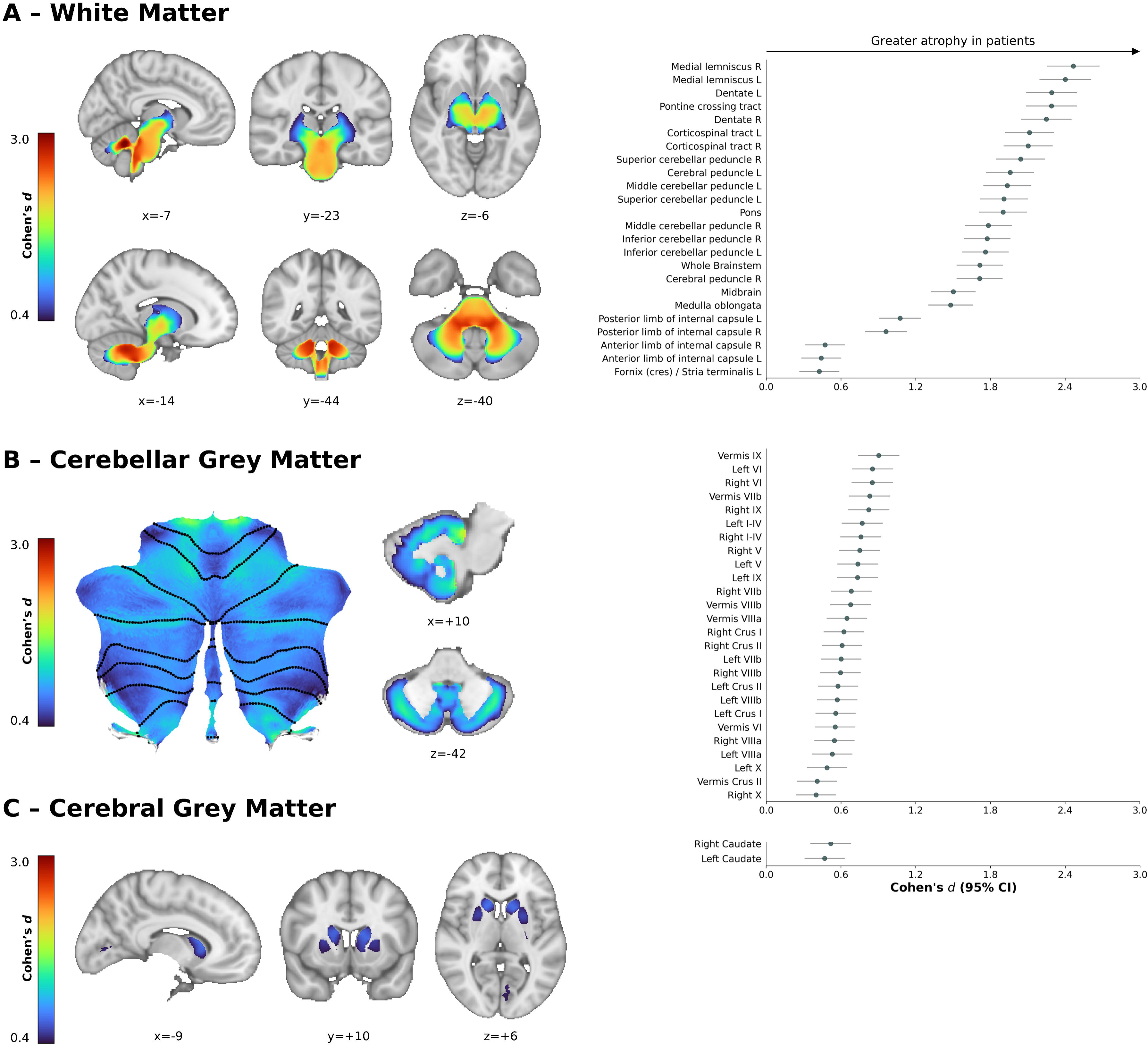
Regions of significantly lower volume (voxel-level FWE-corrected *p* < 0.05) in participants with SCA3 relative to CONT in (A) whole-brain white matter, (B) cerebellar grey matter, and (C) cerebral grey matter. Left: representative slices or cerebellar flatmaps illustrating the areas of significant atrophy in SCA3 subjects. Right: forest plots illustrating regional effects (Cohen’s *d* > 0.4, matching the scale of the brain images); error bars represent the 95% confidence interval (CI). Slice coordinates are in Montreal Neurological Institute (MNI) space.

Analysis of the initial dataset found fundamentally similar results, with a few extra notable sites of significant GM atrophy (Supplementary Figure S2).

### Network Representation of Between-Group Differences

The cerebellar GM results were subsequently mapped onto the MDTB functional regions to identify which functions were most affected by atrophy (Figure 3). From this, three sets of regions emerged based on their relative atrophy level. The highest atrophy was observed in MDTB Regions 5-7 (*d* ≅ 0.7), which are associated with attention, executive function, and language/emotional processing. The next tier of atrophy (*d* ≅ 0.6) was observed in MDTB Regions 1, 2, 8, and 9; the former two represent motor planning and execution while the latter two represent verbal processing. Finally, the weakest atrophy (*d* < 0.5) was observed in regions 3, 4, and 10, which represent a variety of predominantly non-motor skills.

**Figure 3.**
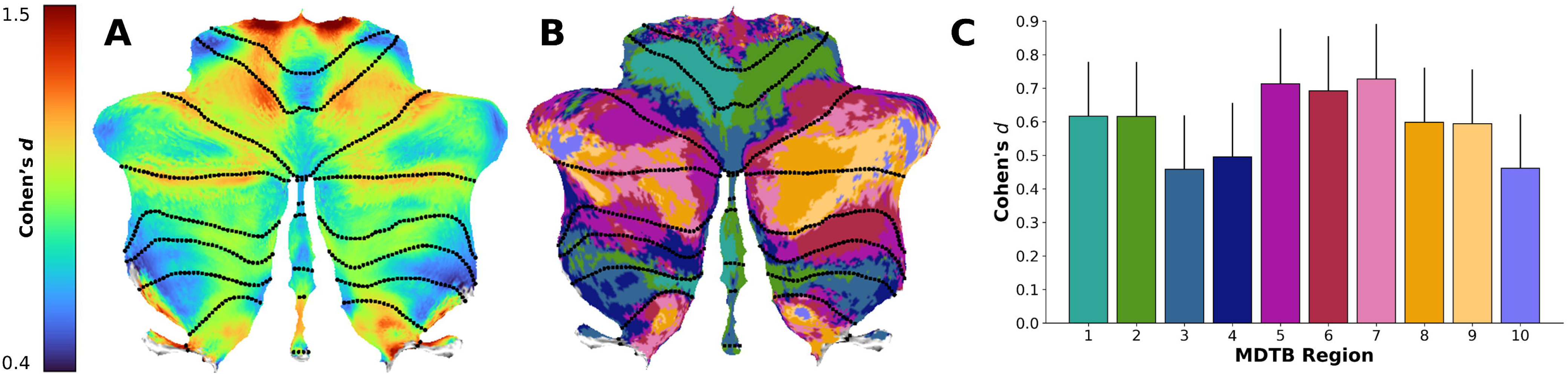
Mapping of structural changes to functional networks in the cerebellum. (A) Cerebellar flatmap of the voxel-level effect size of volume differences in participants with SCA3 vs. CONT in cerebellar grey matter, rescaled from Figure 2B. (B) Cerebellar flatmap of the multi-domain task battery (MDTB) functional atlas from King et al.^22^ (C) A bar chart of the effect sizes by MDTB region; bar height is the mean effect size within each region, while error bars are the positive 95% CI. The colours of the bars in Panel C match the colours of the regions in Panel B.

### Clinical Correlations in SCA3 Participants

Voxelwise correlations between volume and clinical variables were performed to illustrate how atrophy relates to disease duration and intensity (Figure 4). The correlations with both metrics followed similar trends: large negative correlations with volume (0.4 < |*r*| < 0.65) were found in the cerebellar and brainstem WM, while more moderate negative correlations (0.25 < |*r*| < 0.4) were observed throughout much of the cerebellar GM. Correlations with disease duration were subsequently performed in a subset of anatomical ROIs: volumes were z-normalized to the CONT cohort as previously described.^12^ Consistent with the voxelwise findings, the WM and cerebellar GM regions selected showed moderate to strong correlations, while the bilateral caudate – which had the strongest atrophy observed in the cerebral GM – did not (Figure 5).

**Figure 4.**
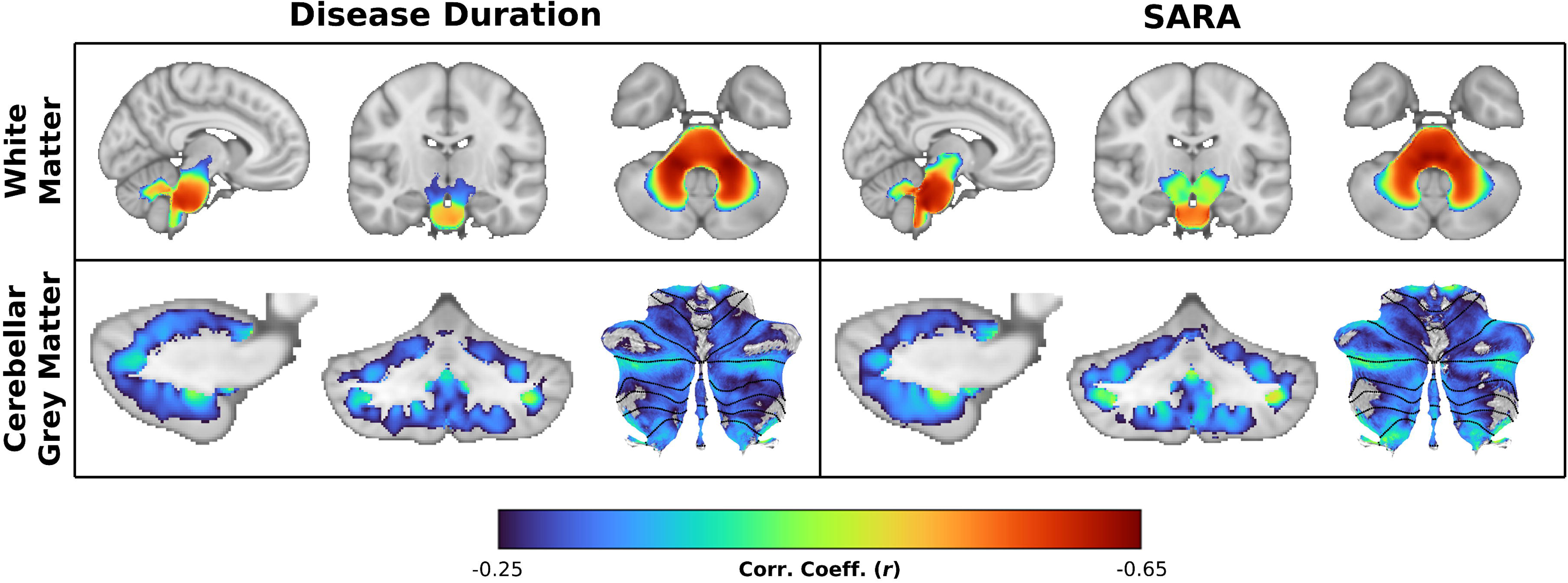
Correlations between volume and disease duration (left) and ataxia severity (SARA score; right) in SCA3 participants (voxel-level FWE-corrected *p* < 0.05). Top row: whole-brain white matter at MNI coordinates (−7,−15,−39). Bottom row: cerebellar grey matter at MNI coordinates: x = +16 and y = −58 with cerebellar flatmap.

**Figure 5.**
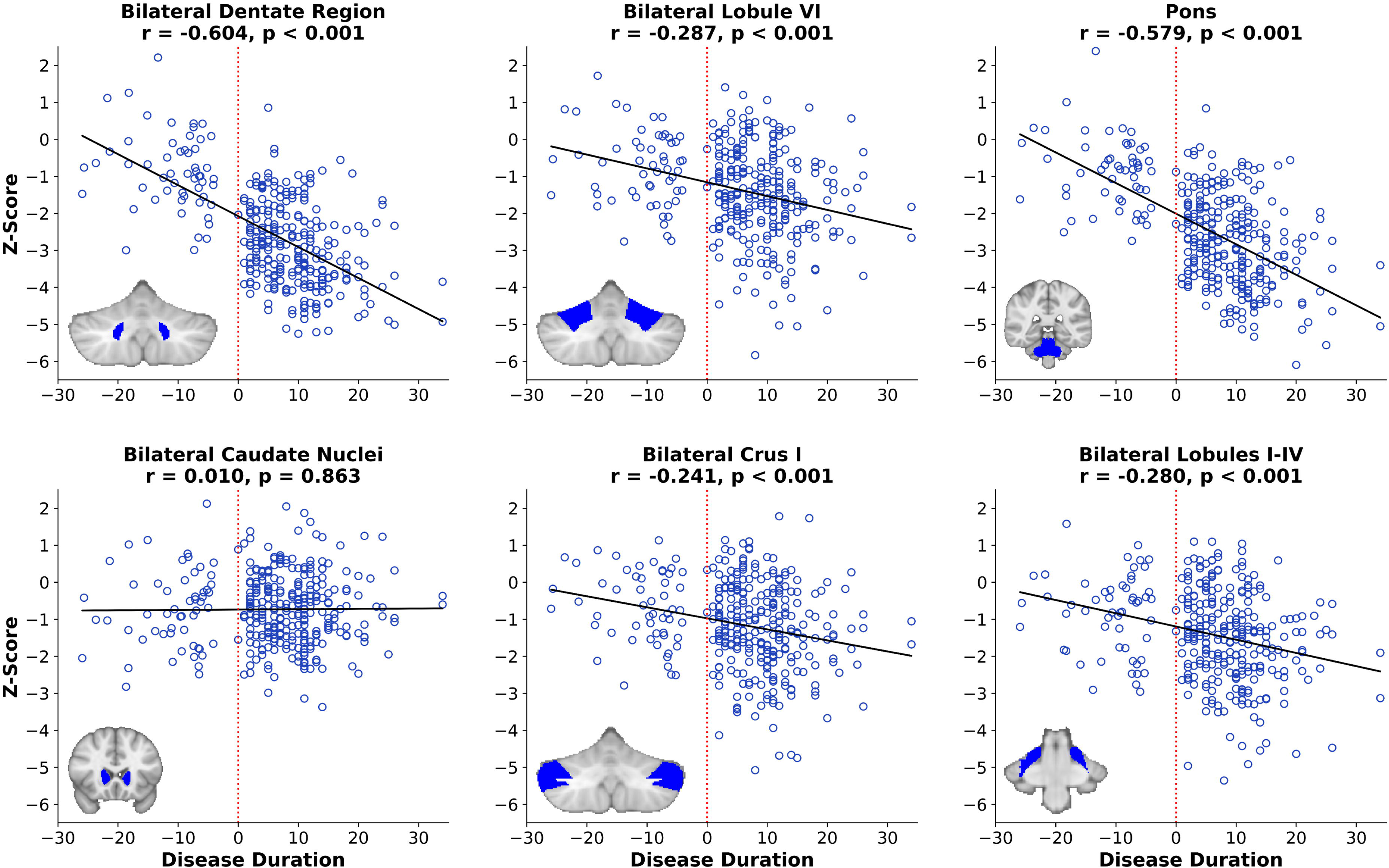
Individual variability within various regions of interest. ROI volume in SCA3 participants is converted to a Z-score within the distribution of CONT participants, then plotted against disease duration in regions with strong disease effects (Cohen’s *d* ≥ 0.8; top row) and moderate disease effects (0.5 ≤ *d* < 0.8; bottom row). Solid line: linear fit between Z and disease duration; dashed line: Duration = 0 line, denoting the separation between pre-ataxic and ataxic SCA3 participants. At the bottom left corner of each plot is an anatomical reference for each ROI’s position in the coronal plane.

Isolated positive correlations were observed in the cerebral WM and GM with respect to disease duration, including small areas of the occipital cortex, thalamus, precuneus, and inferior temporal gyrus(Supplementary Figure S3). No significant correlations were observed between SARA and cerebral GM volume.

Correlations in the initial dataset, which lacked age and site correction, found significance in a variety of new areas of the cerebral GM in particular (Supplementary Figure S4). However, performing the same correlations on the final dataset without correction produced near-identical results; further analysis appears in the Supplementary Results and Supplementary Figure S5.

### Disease Severity Stratification

To characterize the evolution of atrophy over the course of SCA3, the disease cohort was stratified by SARA score (Figure 6). As a broad trend, tissue atrophy becomes more severe as disease severity increases, particularly in the brainstem and cerebellar WM and GM, reflecting the findings of the correlation analysis. In the WM, the spatial extent of the atrophy does not change appreciably across the disease course, however the severity of the atrophy increases markedly with the progression of the disease. In cerebellar GM, pre-ataxic subjects show limited but significant atrophy, predominantly in the anterior regions, which spreads to the rest of the cerebellum and intensifies in the later stages of the disease. In the cerebral GM, significant atrophy is evident in the striatum at all disease stages, with no evidence of spreading or intensification across disease stages; this lack of consistent or progressive involvement of cerebral GM is consistent with the groupwise results.

**Figure 6.**
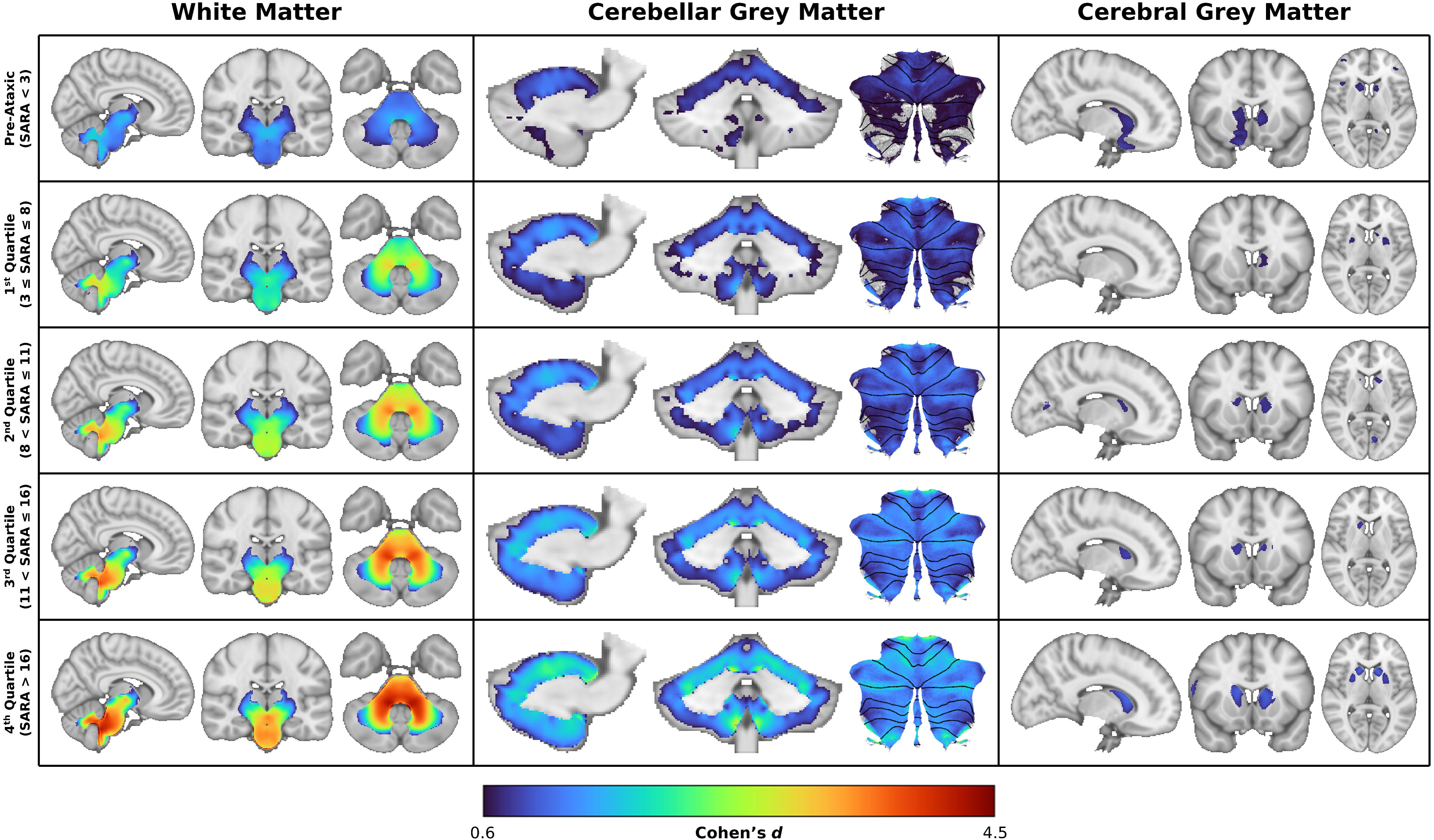
Stratification of the SCA3 cohort based on SARA score. Voxel-level effect size maps (FWE-corrected *p* < 0.05) of each SCA3 subgroup relative to CONT. From top to bottom: Pre-Ataxic (SARA < 3; n = 78), 1^st^ Quartile (3 ≤ SARA ≤ 8; n = 75), 2^nd^ Quartile (8 < SARA ≤ 11; n = 62), 3^rd^ Quartile (11 < SARA ≤ 16; n = 58), 4^th^ Quartile (SARA > 16; n = 60). All images show representative slices or flatmaps of the affected tissue type: white matter (left; MNI coordinates +8,-20,-38), cerebellar grey matter (centre; MNI coordinates x = −13 and y = −48), and cerebral grey matter (right; MNI coordinates +13,+13,+7).

We also observed an unusual significant result in the right frontal-orbital cortex in pre-ataxic subjects, which subsequently disappears in the ataxic subjects. This finding also appears in the groupwise analysis of the initial dataset (Supplementary Figure S2) but nowhere else. To further explore and validate this observation, we present the stratified data at sub-significance thresholds in Supplementary Figure S6.

### Cerebrocerebellar Covariance

The linear models representing cerebrocerebellar covariance in GM, WM, and total volume all produced significant fits, with significant main effects of both *Group and cerebellum*, as well as *cerebellum X Group* interactions. The model thus indicates significant relationships between cerebral and cerebellar volume throughout, but differing magnitudes of these relationships in the CONT and SCA3 groups. Partial regression plots of the cerebrocerebellar relationships, controlled for ICV (Figure 7), show that in all cases, CONT subjects had a steeper slope and a higher *r*^2^ value. In both cohorts, the strongest cerebrocerebellar covariance was observed in the WM. All residual correlations were significant (*p* = 0.009 for SCA3 GM; all others *p* < 0.001). Detailed model statistics are provided in Supplementary Table S2.

**Figure 7.**
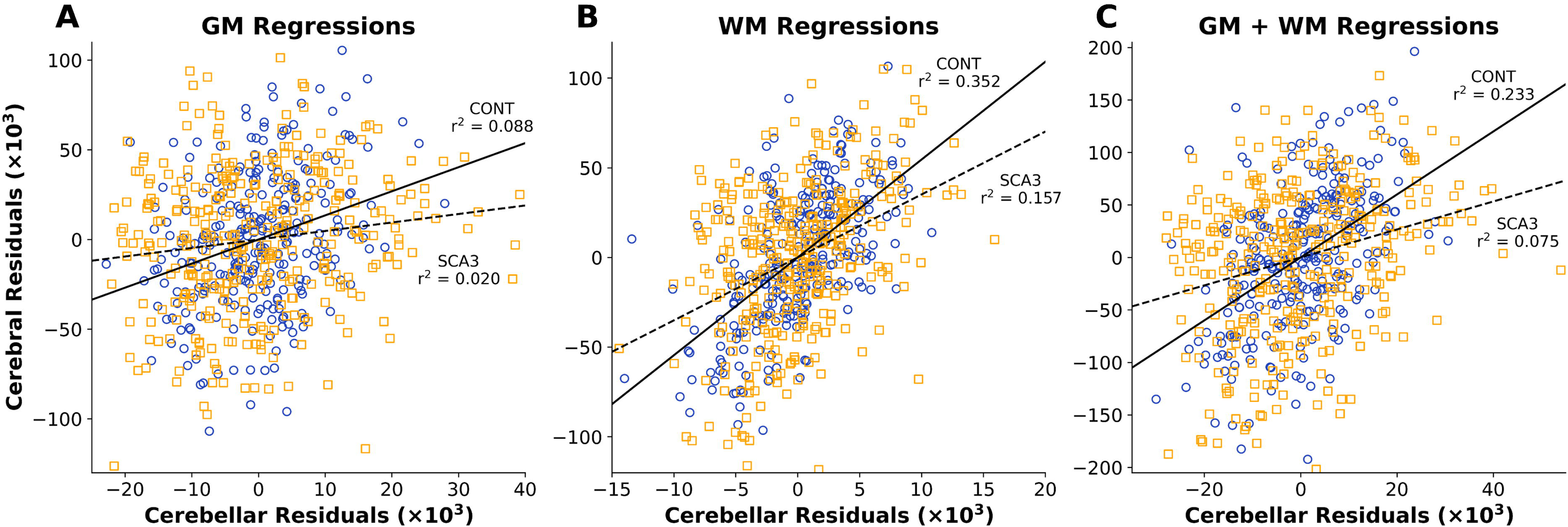
Cerebrocerebellar covariance partial regression plots. Axes represent the residuals from the correlations between ICV and cerebral (vertical) and cerebellar (horizontal) volumes in (A) grey matter, (B) white matter, and (C) both tissues combined. Correlation coefficients for each cohort are labelled on the graph. Blue circles: residual volumes in CONT subjects; orange squares: residual volumes in SCA3 subjects.

## Discussion

In this multi-site MRI study of participants with SCA3 and matched controls, we observed the greatest atrophy in the cerebellar WM and brainstem, with more moderate atrophy occurring across the entirety of the cerebellar GM and parts of the basal ganglia. This pattern remained consistent with disease progression, while correlations with measures of disease severity were strongest in the pons and cerebellar WM. Mapping the cerebellar GM results onto a functional atlas showed that the strongest effects occurred in areas related motor control, cognitive processing, and emotional regulation. Cerebrocerebellar covariance analysis found consistently weaker correlations in SCA3 participants across all tissue types, suggesting that cerebellar atrophy is decoupled from whole-brain, systems-level anatomical changes.

### The Core Atrophic Signature of SCA3

Brainstem and cerebellar WM atrophy was the predominant characteristic of SCA3 observed in this study. This finding is consistent with established literature indicating that WM degradation – particularly in the cerebellum, brainstem, and the tracts connecting them with one another and the striatum^5^ – is a common, early, and progressive manifestation of SCA3, occurring prior to the first onset of motor symptoms.^12,13^ Robust correlations with disease duration and ataxia severity observed in this study are also in line with the established body of longitudinal and cross-sectional studies indicating that atrophy in these core regions worsen as SCA3 progresses.^6,7,9–16,20,21,43,44^

More modest effect sizes of GM atrophy were observed across the whole cerebellar cortex, as well as the bilateral caudate and bilateral putamen nuclei, consistent with previous reports.^6,7,12–14,45^ Here we show that the anterior lobe and superior aspects of the posterior lobe (Lobules I-VI) are impacted first in pre-ataxic individuals, followed by Lobule IX and the remainder of the superior posterior lobe (Lobule VII), then finally Lobules VIIIa/b and X. The relative magnitude of atrophy in each region follows the same pattern across all disease stages. Correlations with disease severity and duration were evident across the cerebellar cortex, although particularly evident in areas neighbouring WM, suggesting the potential for partial volume effects to be driving some of this anatomical variability.

This study indicates that atrophy in the striatum appears to be an early, but non-progressive, feature of SCA3. Significant striatal atrophy was evident in the pre-ataxic stage, but did not reliably correlate with increasing disease severity nor duration. This observation is inconsistent with earlier reports that striatal atrophy is progressive in the pre-ataxic and/or ataxic stages.^6,7,12,14^

### Functional Mapping in the Cerebellum

Mapping our cerebellar GM atrophy results onto a task-based functional map^22^ showed a variable magnitude of atrophy across different functional regions of the cerebellar cortex. Although all regions are impacted, a pattern emerged whereby atrophy was greater in functional regions associated with motor execution and planning, attention, and emotional processing, as compared to regions associated with working memory, sematic/episodic memory, action observation, and saccades. This observation is concordant with the known clinical features of SCA3, including both motor ataxia and a high prevalence of cerebellar cognitive affective syndrome (CCAS)^18,19^, which is defined by abnormalities in executive function, language processing, visuospatial cognition, and affective regulation.^17^

### Inconsistent Evidence of Cerebral and Subcortical Involvement

While there is consistent agreement in the literature and this study about the core regions of SCA3 atrophy, there is less agreement about which regions in the cerebral WM, if any, are affected. In the present study, we found that WM atrophy extended to the internal capsule and peri-thalamic regions. By contrast, using diffusion tensor imaging (DTI), Rezende *et al.*^13^ observed lower fractional anisotropy (FA) relative to controls, implying a loss of WM integrity, across the subcortical WM. Other studies have reported reduced integrity in structures such as the forceps major and minor^21^, inferior and superior longitudinal fasciuli^10,21^, thalamus^14,16^, anterior and posterior thalamic radiations^13,21^, corona radiata^7,13,14^, and corpus callosum^13^ in their respective SCA3 cohorts. These differing findings may be due to the generally greater sensitivity of diffusion MRI metrics to WM microstructural differences as compared to the volumetric approach taken here. Additionally, some studies found positive correlations between disease severity or duration and core SCA3 regions, such as the brainstem and thalamus^45^ or the pons^16^. The reason for this disagreement is less clear and requires further study.

We similarly failed to find robust evidence of consistent cerebral cortical GM atrophy in people with SCA3, even at late disease stages as previously reported.^13^ Notably, however, at reduced, non-significant statistical thresholds, trend-level findings with small effect sizes (i.e., 0.3 ≤ *d* ≤ 0.5) are observed (Supplementary Figure S6), suggesting mild reductions of volume in larger parts of the cerebral cortex. That said, there remains no consistent anatomical pattern, nor clear evidence of progression across the subgroups of our stratified analysis. These observations suggest that, while individuals with SCA3 are as a whole more vulnerable to cerebral degeneration than the general population, there remains extensive individual variability in location and magnitude. Further research will be necessary to determine whether the clinical phenomenology and disease course differ in individuals with versus without cerebral involvement.

### Cerebrocerebellar Correlational Analysis

To further investigate the relationship between cerebral and cerebellar atrophy, we performed a structural covariance analysis and observed a strong covariance between cerebral and cerebellar volume in controls, particularly in the WM. Attenuation of this relationship in individuals with SCA3 suggests that volume loss in the cerebellum occurs relatively independently of any cerebral changes. This result provides further weight to our previous conclusion that the consistent profile of cerebellar volume loss in those with SCA3 is not reflected in cerebral pathology.

### Limitations

Our dataset is rich in T_1_-weighted images and standard clinical measures of ataxia severity (SARA), duration, and genetic load (CAG repeats). Conversely, we lack the cognitive testing or functional MRI data that would allow more definitive insights into the relationship between cerebellar changes and non-motor symptoms such as CCAS. Similarly, more in-depth clinical phenotyping would allow opportunities to unravel the natural heterogeneity of SCA3, including distinguishing the neurobiology of clinical subtypes.^1,4,5^ As noted above, diffusion-weighted MRI would allow for a more sensitive assessment of WM microstructural integrity as compared to our current volume-based techniques. Additionally, while the large SCA3 cohort size in our dataset allows for robust comparisons at various disease severity levels, we recognize that cross-sectional stratification analysis is not a perfect proxy for a true longitudinal study. Finally, while cerebrocerebellar structural covariance analyses provide an indication of the relative coupling of cerebral and cerebellar degeneration, a true assessment of cerebrocerebellar connectivity changes in SCA3 will require functional and/or microstructural imaging approaches.^3^

## Conclusions

In this large, multi-site study, we demonstrate that the core neuroanatomical changes in participants with SCA3 occur in a relatively stable pattern of brainstem and cerebellar atrophy that is progressive in magnitude but not spatial extent, particularly after the onset of ataxia symptoms. This atrophy pattern maps onto functional divisions of the cerebellum that are consistent with the motor and non-motor features of the disease. Finally, we show that this core cerebellar atrophy is not directly mirrored by downstream cerebral changes, and that cerebral volume loss, if present, may be defined by higher individual variability. These findings expand our fundamental understanding of the neuroanatomy of SCA3.

## Acknowledgements

The authors would like to acknowledge the contributions of Victor Gadelha, Hellen Della-Justina, Salmo Raskin, Arnolfo de Carvalho Neto, B.M. and C.W.W. Girard, and Elodie Petit. Several authors of this publication are members of the European Reference Network for Rare Neurological Diseases (ERN-RND).

Some data and/or research tools used in the preparation of this manuscript were obtained from the National Institute of Mental Health (NIMH) Data Archive (NDA). NDA is a collaborative informatics system created by the National Institutes of Health to provide a national resource to support and accelerate research in mental health. This manuscript reflects the views of the authors and may not reflect the opinions or views of the NIH or of the Submitters submitting original data to NDA.

Author J.F. was funded by the Advanced Clinician Scientist Programme (ACCENT), funding code 01EO2107; the German Federal Ministry of Education and Research (BMBF); and as a PI of the iBehave Network, sponsored by the Ministry of Culture and Science of the State of North Rhine-Westphalia. The Center for Magnetic Resonance Research is supported by the National Institute of Biomedical Imaging and Bioengineering (NIBIB) grant P41 EB027061, and the Institutional Center Cores for Advanced Neuroimaging award P30 NS076408 and grant S10 OD017974. Collaborating author P.G. is supported by the National Institute for Health Research, University College London Hospitals Biomedical Research Centre UCLH, and CRN North Thames. Collaborating authors P.G. and H.G.M. work at University College London Hospitals/University College London, which receives a proportion of funding from the Department of Health’s National Institute for Health Research Biomedical Research Centre’s funding scheme. P.G. also received funding from CureSCA3 in support of H.G.M’s work. Author I.H.H. received funding from the Australian National Health and Medical Research Council (Ideas Grant 1184403; Investigator Grant 2026191).

Additional financial support was provided to various authors and sites by the following grants and agencies: EU Joint Programme Neurodegenerative Disease Research (JPND); European Reference Network for Rare Neurological Diseases (ERN-RND); National Ataxia Foundation Clinical Research Consortium grant; National Science Foundation grant #EEC-1460674; National Institute of Health/National Institute of Neurological Disorders and Stroke grants R01NS056307, R01NS080816, and U01 NS104326; Johns Hopkins Institute for Clinical and Translational Research support from NIH National Center for Advancing Translational Sciences (NIH/NCATS) grant UL1TR001079; the Jane Tanger Black Fund for Young-Onset Dementia Research; the Netherlands Organisation for Health Research and Development; Foundation for Science and Technology, Portugal; Medical Research Council, Portugal; Regional Fund for Science and Technology, Azores; Servier; the National Ataxia Foundation; ZonMw grant 733051066; Friedreich’s Ataxia Research Alliance (FARA) grant 92133; São Paulo Research Foundation (FAPESP) grant 2013/07559-3 through CEPID/BRAINN; National Natural Science Foundation of China grant 82071910; Senior Medical Talents Program of Chongqing for Young and Middle-Aged grant 514Z395; Young and Middle-Aged Senior Medical Talents Studio of Chongqing grant 524Z28921; Excellent Young Talent Fund of the First Affiliated Hospital of the Army Medical University grant 2024YQBJ-2; German Research Foundation (DFG) grants DE 2516/1-1 and TI 239/17-1; National Council of Science and Technology of Mexico (CONACYT) grant A1-S-10669; General Directorate of Academic Personnel Affairs Support Program for Research and Technological Innovation Projects (DGAPA-PAPIIT) grant IN208625; the Gossweiler Foundation; NIH Big Data to Knowledge (BD2K) award U54 EB020403; and NIH grants R01MH123163, R01MH121246, and R01MH116147.

For a complete list of ENIGMA-related grant support please see: http://enigma.ini.usc.edu/about-2/funding/.

## Data Availability

READISCA data is available at doi:10.15154/3xeb-fz42. All code and data processing instructions are available at https://github.com/HardingLab/enigma-ataxia. Other ENIGMA collaborator data may be requested for further research purposes through a secondary data use proposal submitted to the ENIGMA-Ataxia working group (http://enigma.ini.usc.edu/ongoing/enigma-ataxia).

