## Supplementary Table 1 for "A Core Pattern of Cerebellar and Brainstem Degeneration and Reduced Cerebrocerebellar Structural Covariance in Spinocerebellar Ataxia Type 3 (SCA3): MRI Volumetrics from ENIGMA-Ataxia"

**Supplementary Tables**

**Supplementary Table S1.** Summary of imaging protocols as reported by each site.

|  | **Aachen^b^** | **Baltimore** | **Campinas** | **Curitiba** | **Essen/Halle** | **Florida^b^** | **Mexico** | **Nijmegen** | **Paris** | **READISCA** | **Tübingen** |
| --- | --- | --- | --- | --- | --- | --- | --- | --- | --- | --- | --- |
| **Scanner** | Siemens Prisma | Philips Intera | Philips Achieva | Siemens Skyra | Siemens Biograph | Philips Achieva | Philips | Siemens Avanto | Siemens Trio | Siemens Prisma/ Skyra^b^ | Siemens Skyra |
| **Field (T)** | 3 | 3 | 3 | 3 | 3 | 3 | 3 | 1.5 | 3 | 3 | 3 |
| **Head Coil Channels** | 64 | 8 | 8 | 16 | 16 | 32 | 32 | --^a^ | 32 | 32 | 32 |
| **Sequence** | MPRAGE | MPRAGE | SPGR | MPRAGE | MPRAGE | --^a^ | FFE | --^a^ | MPRAGE | MPRAGE | MPRAGE |
| **TR (ms)** | 2500 | 10.3136 | 7 | 2530 | 2530 | 8.2 | 8 | 1900 | 2530 | 2400 | 2300 |
| **TE (ms)** | 4.37 | 6 | 3.201 | 3.36 | 3.26 | 3.7 | 3.7 | 2.5 | 3.65 | 2.22 | 2.32 |
| **TI (ms)** | 1100 | --^a^ | --^a^ | 1100 | 1100 | --^a^ | 900 | 1100 | 900 | 1000 | 900 |
| **Flip Angle (°)** | 7 | 8 | 8 | 7 | 7 | 8 | 25 | 15 | 9 | 8 | 8 |
| **Plane** | Sagittal | Axial | Sagittal | Sagittal | Sagittal | Sagittal | Sagittal | --^a^ | Sagittal | Sagittal | Sagittal |
| **Slices** | 192 | Variable | 180 | 176 | 176 | 170 | 176 | 176 | 160 | 208 | 192 |
| **Field of View** | 256 × 256 | 256 × 256 | 240 × 240 | 256 × 256 | 256 × 256 | 240 × 240 | 256 × 256 | 256 × 256 | 256 × 256 | 320 ×320 | 230 × 230 |
| **Voxel Size (mm, X × Y × Z)** | 1 × 1 × 1 | 0.828125 × 0.828125 × 1.10 | 1 × 1 × 1 | 1 × 1 × 1 | 1 × 1 × 1 | 1 × 1 × 1 | 1 × 1 × 1 | 0.9 × 0.9 × 1 | 1 × 1 × 1 | 0.8 × 0.8 × 0.8 | 0.9 × 0.9 × 0.9 |

FFE: fast field-echo; IR-3D GRE: 3D inversion recovery gradient echo; MPRAGE: magnetization-prepared rapid gradient-echo; SPGR: spoiled gradient echo; TE: echo time; TFE: turbo field-echo; TI: inversion time; TR: repetition time
^a^ Information not provided.
^b^ Site or scanner used for initial dataset only.

Detailed data unavailable from Chongqing, ESMI, and EUROSCA.

**Supplementary Table S2.** Linear model statistics from cerebrocerebellar covariance analysis.

|  | Model Fit | | | *cerebellum* | | *Group* | | *ICV* | | *cerebellum* × *Group* | |
| --- | --- | --- | --- | --- | --- | --- | --- | --- | --- | --- | --- |
|  | *F*_4,610_ | *p* | Adj. *R*^2^ | *t* | *p* | *t* | *p* | *t* | *p* | *t* | *p* |
| GM only | 142.8 | < 0.001 | 0.480 | 5.153 | < 0.001 | 2.277 | 0.023 | 16.65 | < 0.001 | -2.821 | 0.005 |
| WM only | 239.1 | < 0.001 | 0.608 | 11.58 | < 0.001 | 4.203 | < 0.001 | 13.75 | < 0.001 | -2.835 | 0.005 |
| GM + WM | 231.2 | < 0.001 | 0.600 | 8.819 | < 0.001 | 3.958 | < 0.001 | 17.51 | < 0.001 | -3.791 | < 0.001 |

Adj. *R*^2^: Adjusted *R*^2^; ICV: intracranial volume; GM: grey matter; WM: white matter

**Supplementary Figures**

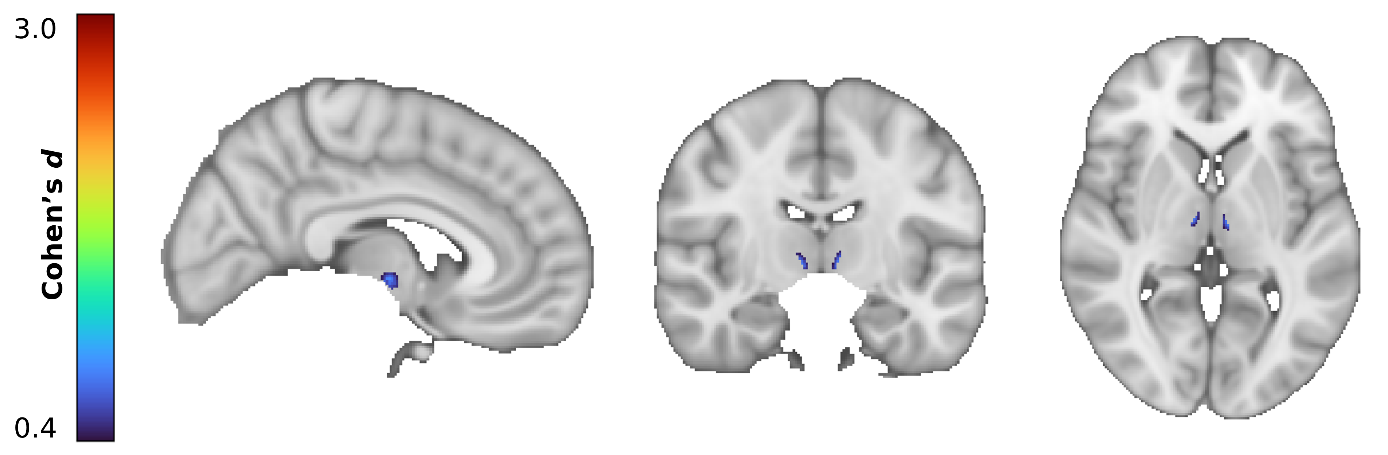

**Supplementary Figure S1.** A small region of the bilateral thalamus which was found to have greater volume in the SCA3 cohort compared to the CONT cohort (voxel-level FWE-corrected p < 0.05). MNI coordinates of slices: (+6,-13,+3).

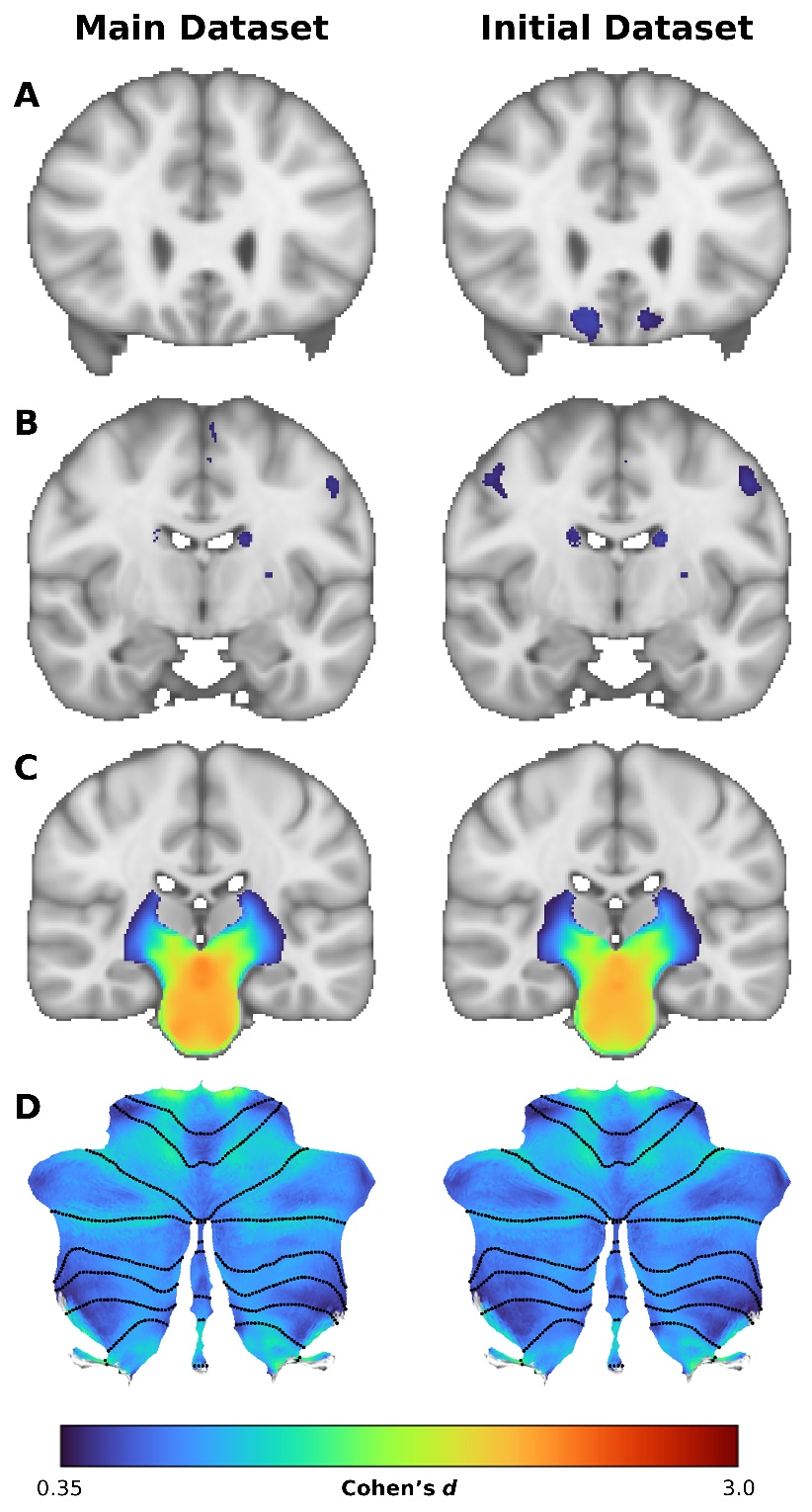

**Supplementary Figure S2**. Images showing the differences between findings from the main dataset (left) and the initial dataset (right). (A) Cerebral grey matter at y = +26, showing the addition of the bilateral frontal orbital and subcallosal cortices. (B) Cerebral grey matter at y = -8, showing the addition of the right precentral gyrus. (C) White matter at y = -23, reflecting a slight decrease in effect size. (D) Cerebellar grey matter flatmap, reflecting a slight decrease in effect size.

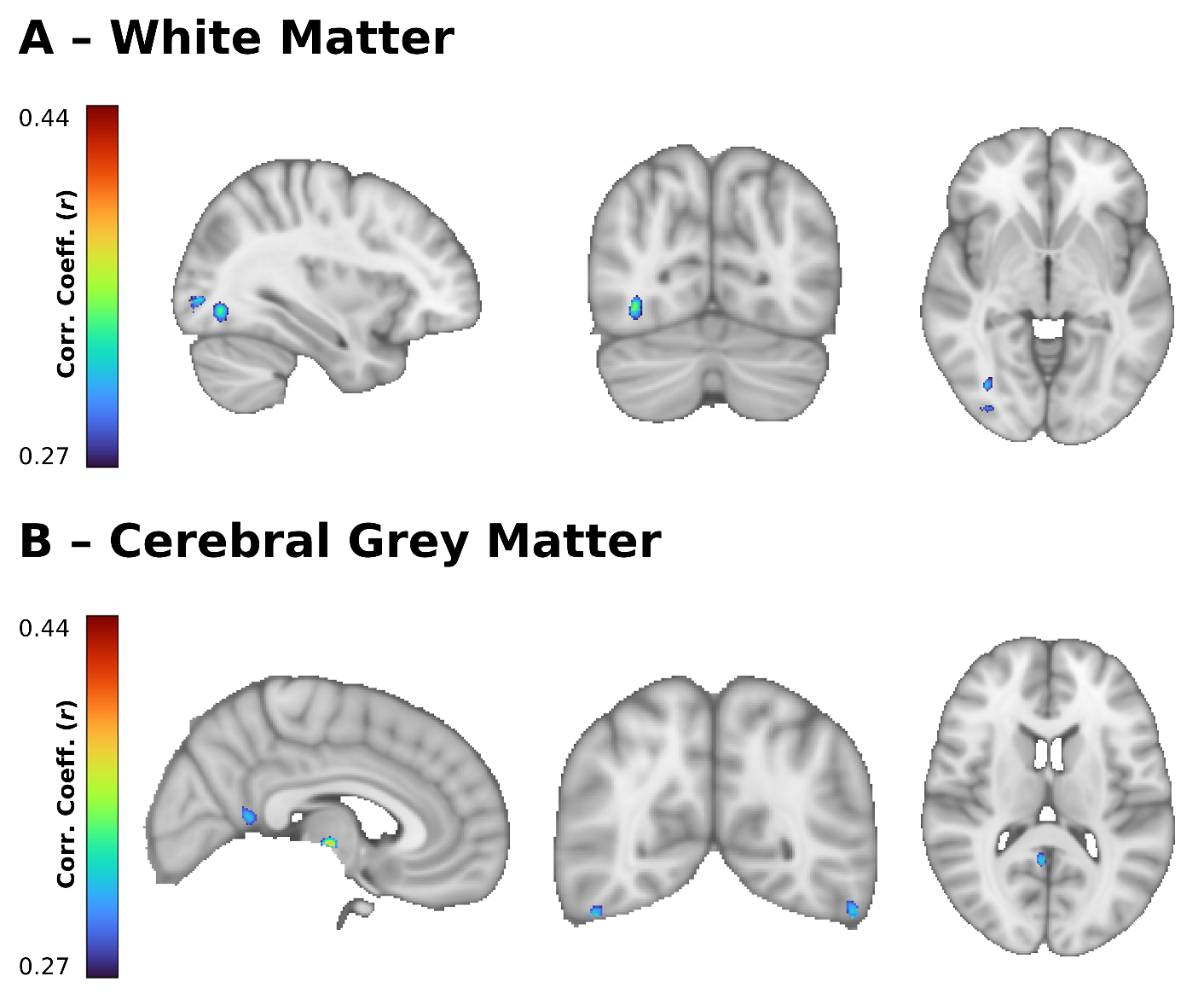

**Supplementary Figure S3.** Positive correlations of volume with disease duration in SCA3 participants. (A) White matter correlations taken at MNI coordinates (+34,+21,-1); (B) Cerebral grey matter correlations taken at MNI coordinates (+5,-59,+13).

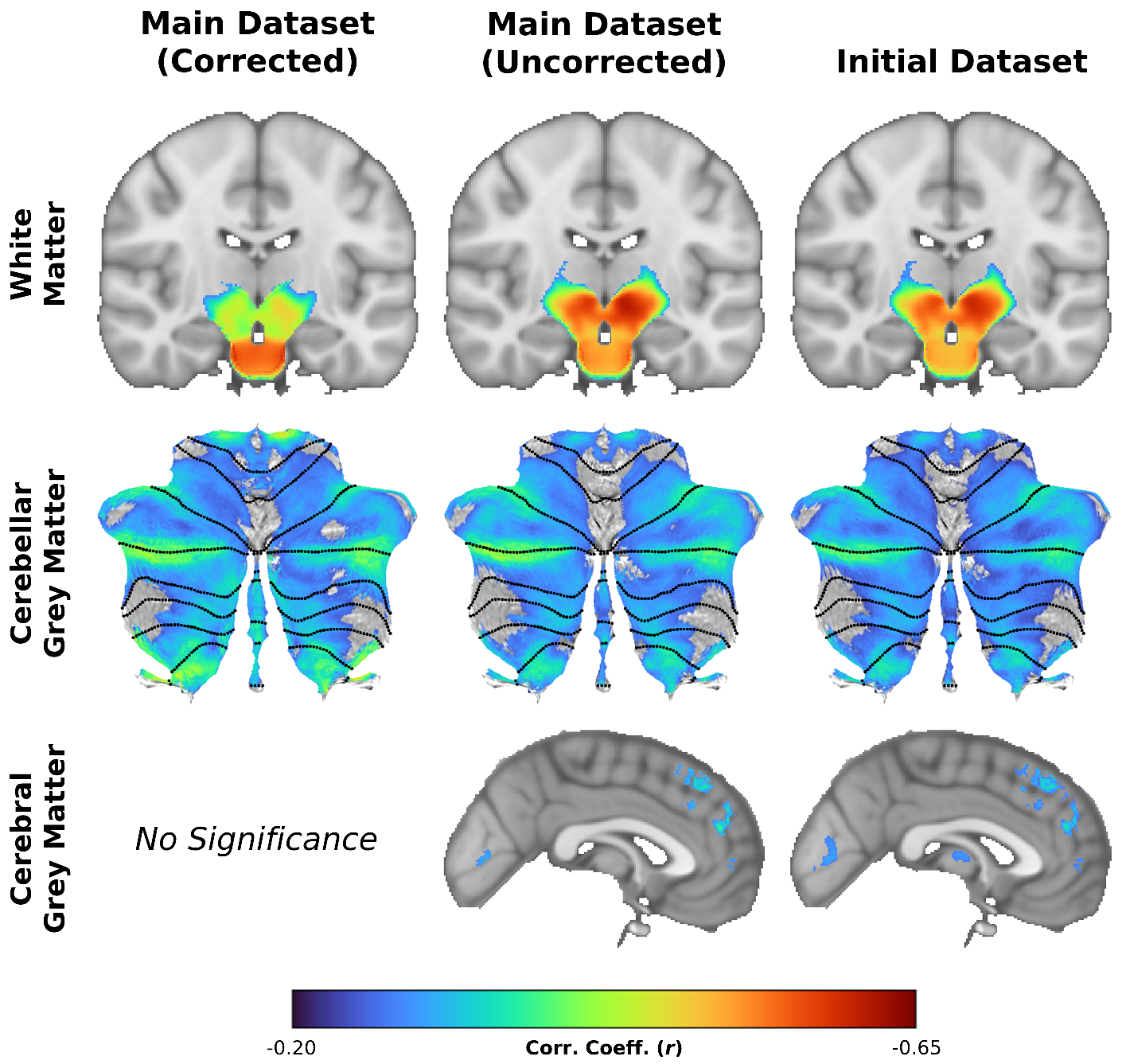
**Supplementary Figure S4.** Images showing differing correlations between SARA and volume using the age- and site-corrected main dataset (left), an uncorrected version of the main dataset (middle), and the initial dataset (right). Top row: whole-brain white matter at y = -15. Middle row: cerebellar grey matter flatmap. Bottom row: cerebral grey matter at x = -1.

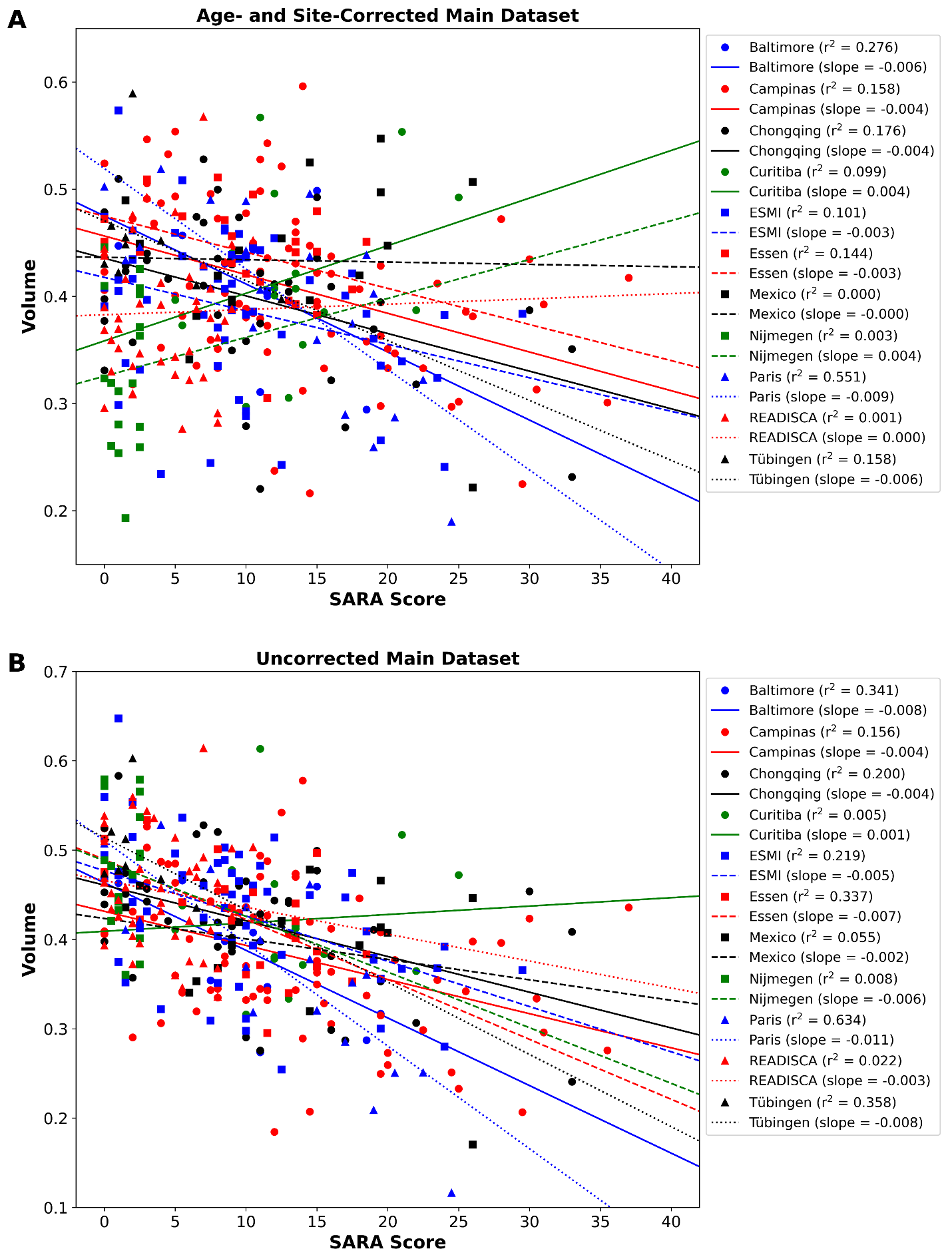

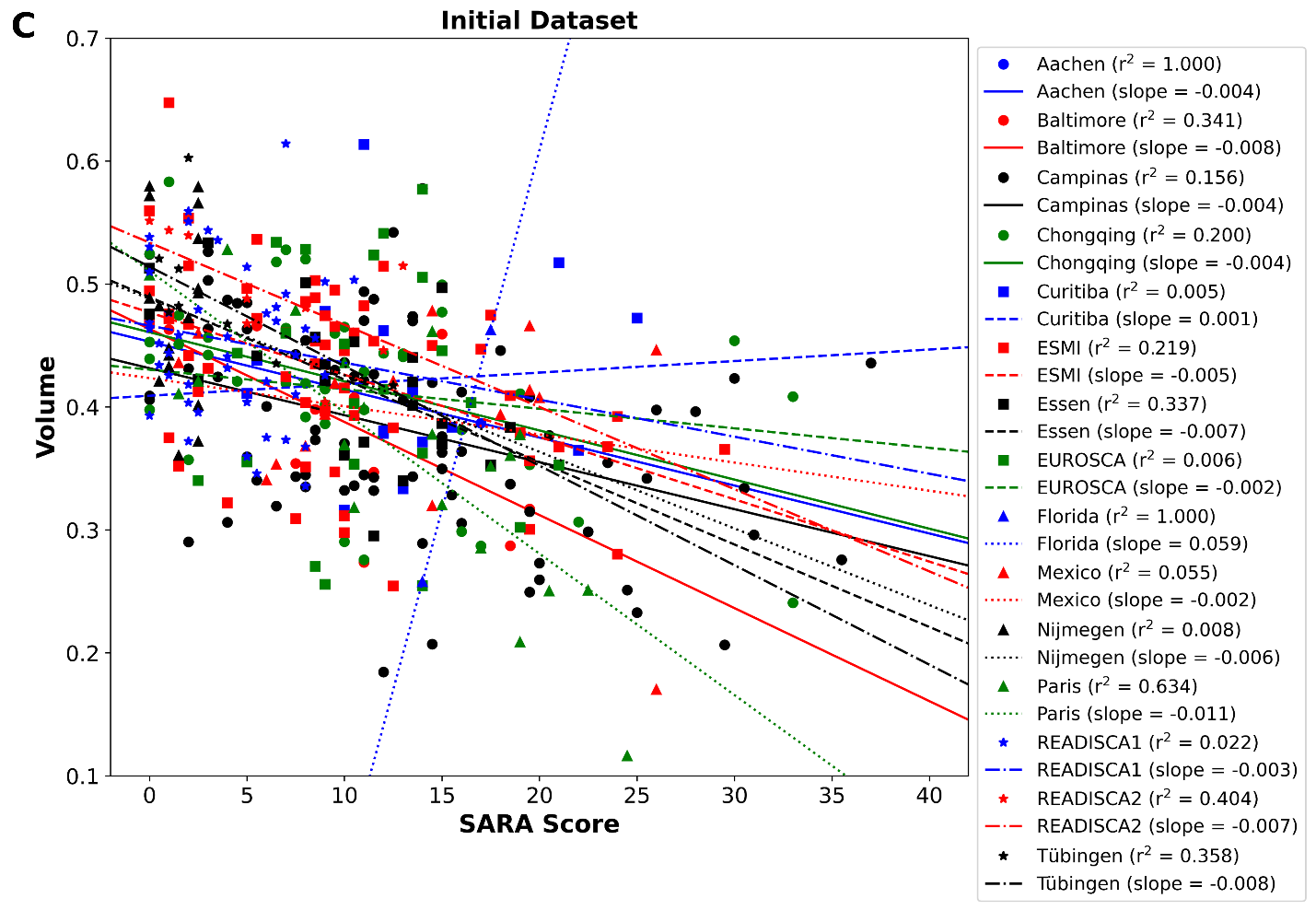

**Supplementary Figure S5.** Correlations between SARA score and volume in a sample of 125 voxels in the cerebral grey matter, with per-site correlation values (A) using the age- and site-corrected main dataset; (B) using the main dataset without age- and site-correction; and (C) using the initial dataset.

**Commentary:** Supplementary Figure S4 illustrates the differences between the main (left) and initial dataset (right). While the distinction in the significance maps was small for the WM and cerebellar GM, the large and scattershot nature of the newfound significances in the cerebral GM gave us pause. Given that the difference between the two datasets was very small in terms of real numbers (339 vs. 376, respectively), we then decided to examine the main dataset without age and site correction to determine whether this was an artefact of our inability to apply said corrections due to lack of sufficient controls. The results from the uncorrected main dataset (middle column) were nearly identical to those in the initial dataset, confirming that the lack of correction was the culprit.

Next, we extracted volume data from a 5x5x5 mm cube of voxels in one of the newly-significant regions of the cerebral GM containing a local maximum, centred on MNI coordinates (-3,+47,+24), and plotted that against SARA score (Supplementary Figure S5). The purpose of this was to determine whether the primary driver of these differences was age or site correction. If the differences were primarily site correction driven, then the lines of best fit would change position but not slope nor $r^{2}$, as site correction is performed by subtracting the mean control image, whereas age correction is performed with a linear model that would affect this relationship. As we observed the latter, we find it likely that both age and site correction were important to removing superfluous relationships, thus affirming our decision to exclude the uncorrectable data (see Figure 1).

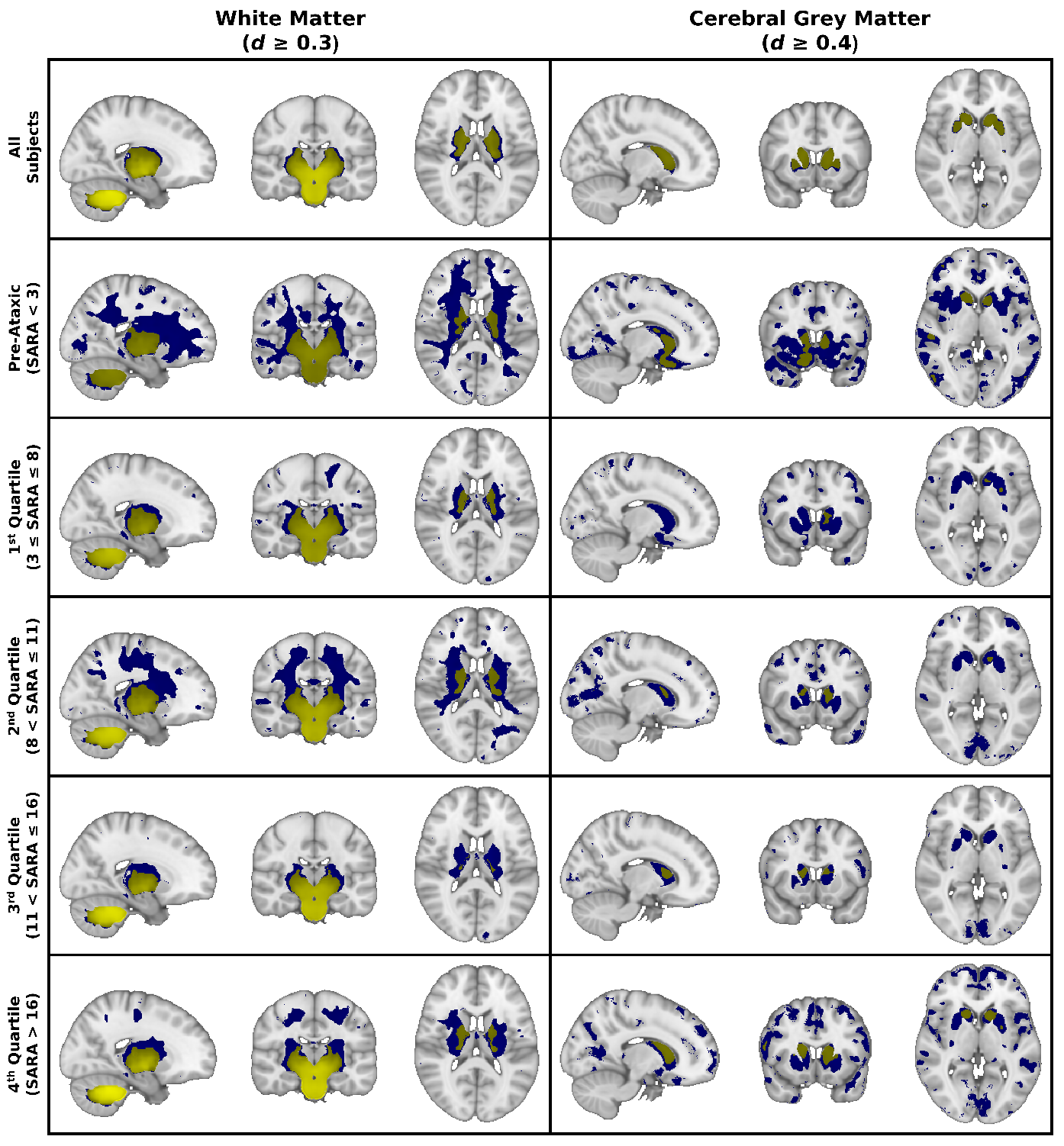

**Supplementary Figure S6.** Observable trends in non-significant clusters of voxels in the staging analysis of white matter (left) and cerebral grey matter (right). Rows, in descending order, depict the full groupwise comparison, followed by the pre-ataxic subjects, followed by the four quartiles of SARA in sequence. (See Figure 6 for more details.) Yellow voxels are significant at *p*_FWE_ < 0.05; blue voxels meet the effect size threshold listed at the top of the column. MNI coordinates for WM comparison are (+20, -18, +14); for the GM comparison (+14, +12, +2).
